# *In vivo* binding by Arabidopsis SPLICING FACTOR 1 shifts 3′ splice site choice, regulating circadian rhythms and immunity in plants

**DOI:** 10.64898/2025.12.17.693997

**Authors:** Yamila Carla Agrofoglio, María José Iglesias, María José de Leone, Carlos Esteban Hernando, Martin Lewinski, Sol Belén Torres, Giuliana Contino, Marcelo Yanovsky, Dorothee Staiger, Julieta Lisa Mateos

**Affiliations:** Instituto de Fisiología, Biología Molecular y Neurociencias (IFIBYNE-UBA-CONICET) and Facultad de Ciencias Exactas y Naturales, Universidad de Buenos Aires, Ciudad Universitaria, Buenos Aires, Argentina; Fundación Instituto Leloir, Instituto de Investigaciones Bioquímicas de Buenos Aires–Consejo Nacional de Investigaciones Científicas y Técnicas (CONICET), C1405BWE Buenos Aires, Argentina; RNA Biology and Molecular Physiology, Faculty of Biology, Bielefeld University, 33615 Bielefeld, Germany

## Abstract

Alternative splicing expands proteome diversity and enables phenotypic plasticity across eukaryotes. In plants, mutations in spliceosomal components impair development and stress responses, but the molecular mechanisms remain unclear. Here, we define the molecular function of SPLICING FACTOR1 (AtSF1) in *Arabidopsis thaliana* using individual-nucleotide resolution UV crosslinking and immunoprecipitation (iCLIP) combined with RNA sequencing of *sf1* mutants. We identify the *in vivo* branch point sequences bound by AtSF1 and delineate its RNA-binding landscape, revealing pervasive splicing defects dominated by aberrant 3′ splice site selection. Structural comparison with human SF1 indicates that AtSF1 retains branch point recognition capacity but features a distinct domain organization, including a restructured C-terminal region absent in metazoans, suggesting a divergent RNA-binding mode that evolved to meet plant-specific splicing demands. AtSF1 targets are enriched for core circadian clock and defense genes, consistent with the long-period phenotype and immune-compromised phenotypes of *sf1* mutants. Together, these findings establish that AtSF1 orchestrates alternative 3′ splice site choice through intron binding and branch point recognition, coupling RNA processing with circadian and immune regulation in plants.

**Significance:** Pre-mRNA splicing is a fundamental process that shapes gene expression and proteome diversity, yet how it integrates with physiological pathways in plants remains poorly understood. Our study identifies the spliceosomal component SPLICING FACTOR1 (AtSF1) as a central modulator of alternative 3′ splice site choice in *Arabidopsis thaliana*. By defining branch point sequences and direct RNA targets of AtSF1 *in vivo*, we reveal its dual regulatory role in circadian timing and immune responses. Comparative analysis with human SF1 uncovers a distinct domain architecture in the plant homolog, suggesting an alternative RNA-binding mode that evolved to meet plant-specific demands. These findings illuminate how conserved splicing machinery was molecularly adapted in the plant lineage to coordinate RNA processing with environmental and developmental cues.

## INTRODUCTION

Pre-mRNA splicing is an essential step in eukaryotic gene expression, removing introns that are defined by short conserved sequence motifs (the 5’ and 3’splice sites) and joining exons to generate mature mRNAs. Many genes give rise to two or more mRNA isoforms, depending on which regions are treated as exons during RNA processing, a process known as alternative splicing (AS). In that way, AS enriches protein diversity and phenotypic traits. It has been proposed that AS is a source of phenotypic plasticity and contributes to evolution in eukaryotes (1). Splicing is catalysed by the spliceosome, a dynamic ribonucleoprotein complex composed of five small nuclear ribonucleoprotein particles (snRNPs) and more than 200 accessory proteins (2–4). Early spliceosome assembly is initiated by the recognition of the 5’splice site (5’ss) by U1 snRNP and recruitment of SPLICING FACTOR1 (SF1), which binds to the branch point sequence (BPS) and facilitates association of the two U2 small nuclear ribonucleoprotein auxiliary factors U2AF65 and U2AF35 to the polypyrimidine tract and the AG dinucleotide at the 3’ss, respectively. This is known as the spliceosomal early complex E. In particular, interaction of SF1 with the BPS is thought to facilitate base pairing of the U2 snRNA. This bulges out the adenosine residue (5). In the presence of ATP, the E complex is remodelled to the A complex with U2 snRNP associated with BPS (3).

The branch point is a critical *cis*-element for splice site selection and intron removal. The branch point adenosine executes a nucleophilic attack on the 5’ss, liberating the upstream exon and generating an intermediate lariat structure from the intron. In the second step of the splicing process, the upstream exons execute a nucleophilic attack on the 3’ss, joining the exons and liberating the lariat which is subsequently hydrolysed by the RNA debranching enzyme DBR1 (6). Debranching of the lariat results in a linear RNA that is further degraded by exonucleases.

In animal and yeast, BPS recognition has been mapped through biochemical and high-resolution binding assays (7, 8). In plants, however, the BPS consensus has been less well defined. Bioinformatic analysis of Arabidopsis introns proposed a WWCUR**A**W motif (9, 10), while lariat sequencing in *DEBRANCHING ENZYME 1* mutants identified an uracil-rich context surrounding the conserved adenosine (11). Notably, ∼20% of introns contain multiple candidate branch point (11), underscoring the complexity of BPS usage in plants.

SF1 is a conserved spliceosomal component. In mammals and yeast, SF1 contains a K Homology (KH) domain located upstream of the Quaking homology (QUA) 2 region, two zinc knuckle domains, and a C-terminal proline-rich region, where the KH and QUA2 domains directly recognize the BPS (12–17). In plants, the SF1 ortholog retains these domains but additionally uniquely carries an RNA recognition motif (RRM) upstream of the proline-rich region (18, 19). This RRM contains the conserved Ribonucleoprotein 1 (RNP1) and RNP2 motifs and is required for full complementation of the early flowering phenotype of *sf1* mutant in Arabidopsis (20), yet its precise molecular contribution to RNA binding remains unresolved.

Beyond their role in splicing, splicing factors in plants often participate in multiple biological pathways, from regulation of the circadian clock (21–26) to responses to abiotic and biotic stress (21, 23–25, 27–29). In Arabidopsis, the circadian clock relies on interconnected transcriptional–translational feedback loops to maintain a ∼24-hour rhythm, aligning plant physiology with the day–night cycle (30). Key components such as CIRCADIAN CLOCK ASSOCIATED1, LATE ELONGATED HYPOCOTLY, and TIMING OF CAB EXPRESSION 1 (TOC1), along with pseudoresponse regulators (PRRs) like PRR7 and PRR9 and evening complex members (LUX ARRHYTHMO, EARLY FLOWERING 3 and 4), govern this precise timing through reciprocal regulation (31). Rhythmically expressed clock proteins convey rhythmicity on output genes. While these transcriptional feedback loops are well characterized, AS has recently been recognized as a crucial additional layer influencing circadian rhythm (23–25, 32, 33). On the other hand, we and others have previously showed that AS of key defence genes such as intracellular immune receptors including the nucleotide-binding leucine-rich repeat (TIR-NB-LRR) protein family is highly dynamic during bacterial infection and plays a role in plant defence (21, 34–36). Unlike in yeast, worm, fruit fly, mice and humans- where SF1 is essential and loss-of-function mutations are lethal (14, 37–39), Arabidopsis *sf1* mutants are viable, albeit with pleiotropic phenotypes including dwarfism, early flowering, and abscisic acid hypersensitivity (40). This unique viability provides a powerful opportunity to dissect AtSF1 function in splicing within a multicellular organism addressing at the same time its impact on plant biology.

Here, we integrated individual-nucleotide resolution UV crosslinking and immunoprecipitation (iCLIP) with transcriptome profiling to defined the AtSF1 RNA-biding landscape and its role in splicing regulation. We identified AtSF1-bound branch point motifs *in vivo* and demonstrated that AtSF1 controls in 3’ss selection. In *sf1* mutants, alternative 3′ss positioned upstream of canonical sites are preferentially used, resulting in longer transcript isoforms. Our analysis revealed widespread splicing defects affecting circadian clock and defence-related genes. Consistent with these molecular alterations, *sf1* mutants displayed a long circadian period and increased susceptibility to bacterial infection. Overall, our findings provide mechanistic insights into how AtSF1 shapes splicing though branch point recognition and establish its central role linking RNA processing to essential biological processes including clock regulation and pathogen defence.

## RESULTS

### AtSF1 preferentially binds introns *in planta*

Although SF1 has been thoroughly characterized in yeast and humans (12), Arabidopsis AtSF1 displays a unique domain architecture (40) that could influence RNA binding dynamics. To elucidate AtSF1’s molecular interactions and its role in splicing, we performed transcriptome-wide mapping of RNA targets using iCLIP (41, 42). To this end, we generated transgenic Arabidopsis lines expressing *GFP:AtSF1* in *sf1-4* T-DNA insertion mutant background (43) alongside *GFP*-only lines used as negative controls. The *GFP:AtSF1* transgene complemented the small rosette phenotype of the *sf1-4* mutant, confirming that the fusion protein is biologically active (Figure 1A). Following UV crosslinking and immunoprecipitation (IP) with GFP-trap^®^ beads, RNA-GFP:AtSF1 complexes were recovered as visualized in a gel after radioactive labelling of the RNAs (Figure 1B). Treatments of IP fractions with RNase I abolished the radioactive signal, verifying that RNAs were co-precipitating (Figure 1B). iCLIP libraries from four independent GFP:AtSF1 replicates and three GFP*-*only controls were sequenced (Figure S1), with read statistics detailed in Table S1.

**Figure 1.**
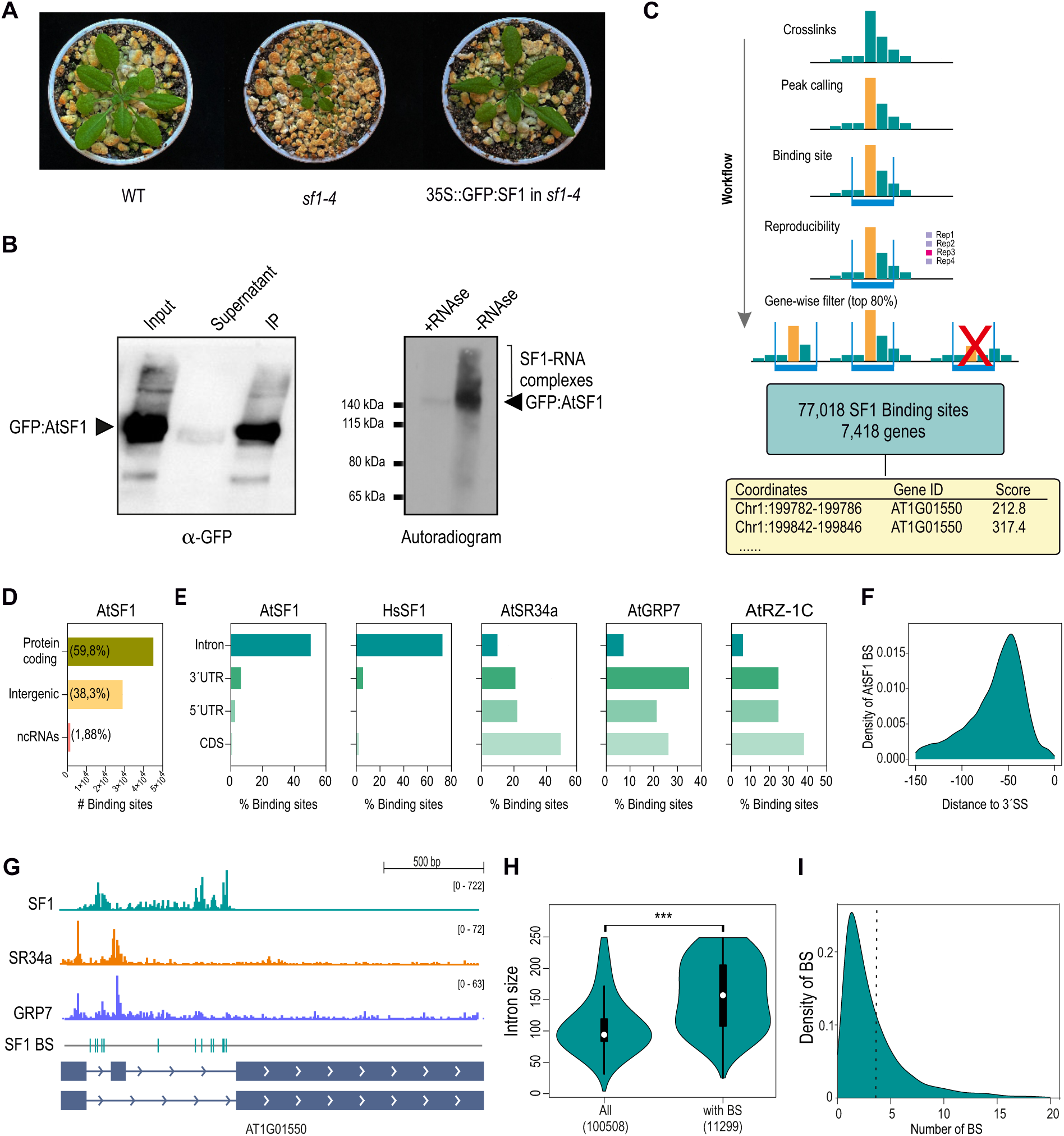
Genome-wide identification and characterization of AtSF1 RNA-binding sites by iCLIP. (A) Rosette phenotype of wild type, *sf1-4* and *sf1-4* plants complemented with SF1:GFP used for iCLIP. Pictures were taken 25 days after sowing (B) Left: Immunoblot analysis of the IP performed in iCLIP conditions and developed with an anti-GFP antibody. Right: Autoradiogram of AtSF1-GFP protein–RNA complexes. After ultraviolet crosslinking, cell lysates were subjected to immunoprecipitation with GFP Trap beads. RNAs were radioactively labelled, and the complexes were separated by denaturing gel electrophoresis. Treatment of the lysate with RNase I (+ RNase) indicates the size of the precipitated proteins. The region above the fusion protein containing the co-precipitated RNAs used for library preparation is indicated. (C) Schematic workflow of processing iCLIP reads and calling SF1 binding sites. The total number of BSs and target genes are indicated. (D) Gene classes targeted by AtSF1 based on iCLIP. (E) Comparison of transcript regions of protein-coding genes targeted by different splicing-related proteins: AtSF1 (this study), HsSF1 (ref), AtRZ-1C (ref), AtGRP7 (ref), AtSR34a (ref). All data was based on iCLIP experiments from different labs. (F) Genome browser tracks of SF1, SR34a and AtGRP7 iCLIP in shared target comparing binding patterns through read density. (G) Intron size distribution of SF1 binding sites. P-Val was determined by Mann-Whitney test. (H) Density of SF1 binding sites in introns.

Uniquely mapped reads were processed using PureCLIP for peak calling (44) (Figure 1C, Figure S2), resulting in 1,132,22 crosslink peaks from which 77,018 binding sites (BSs) were identified (Table S2) using the *BindingSiteFinder* R package (45). The mean of the BS width resulted in 5 nucleotides (Figure S3A). All four replicates displayed reproducible log-normal crosslinks distribution (Figure S3B). Only the top 70% scoring BSs per transcript were retained. Cross-referencing to AtRTD3 gene models (46) identified AtSF1 binding in 7,418 genes, with <0.005% of the total BSs mapping to organellar genes (Figure 1C, Table S3). Most AtSF1 BS overlapped with protein-coding genes (59.8%, Figure 1D), while only ∼2% mapped to noncoding RNAs. Within coding genes, AtSF1 binding was highly enriched in introns versus other regions: <1% of the BSs mapped to coding regions (CDS), 6.3% to 3’UTRs and 2.3% to 5’UTRs (Figure 1E). Notably, this intronic preference is conserved in Human SF1 (HsSF1) (8) (Figure 1E), indicating that SF1 binding is conserved across kingdoms. Yet, this contrasts with the CDS or UTR bias found for other plant splicing factors whose direct RNA targets have been characterized by iCLIP (Figure 1E). Serine-Arginine-rich proteins and hnRNP-like protein AtRZ-1c preferentially bind to the CDS (47–49), while the hnRNP-like *At*GRP7 predominantly binds to the UTRs (50).

AtSF1 crosslinks span entire introns but BSs concentrate towards the 3’ end (Figure 1F-G). Arabidopsis introns are considerably shorter than mammalian ones–median length of 160-170 nucleotides compared to several kilobases in human genes (51–53). Though 85% of Arabidopsis introns fall below 250 nucleotides (∼100 bases average length), AtSF1 binds preferentially to longer introns (∼150 bases, *p-*value < 2.2 e-16) (Figure 1H), with most introns averaging three and some introns (∼300) exceeding fifteen (Figure 1I).

### AtSF1 binding defines plant BPS but is not positioned at the branch point

While SF1 binding preferences have been well studied in yeast and humans (7, 8, 54), little experimental evidence exists for plants. To address this, we performed a *de novo* motif discovery around the AtSF1 BSs defined above using a 30-nucleotide window. Motif enrichment analysis revealed two predominant intronic motifs (Figure 2A): a U-rich tract (motif 1), present in 86% of BSs (Figure 2A), and the CUR**A**Y sequence (motif 2), which closely matches the predicted plant BPS determined bioinformatically in Arabidopsis (9–11), indicating that this motif corresponds to the BPS preferentially recognized by AtSF1. Remarkably, motif 1 is localized almost directly at the AtSF1 BS, whereas motif 2 is consistently found ∼10–25 nucleotides from the center of the AtSF1 BS (Figure 2B). Alignment of AtSF1 BSs with lariat-defined branch points (11), revealed distinct enrichment peaks at approximately 9 nt and 30 nt upstream of the BPS (Figure 2C). Collectively, these data demonstrate that AtSF1 binds near, but not exactly at, the branch point, with occupancy adjacent to BPS-like motifs. Our iCLIP and motif analyses provide in vivo evidence for BPS motif enrichment at AtSF1 binding sites in Arabidopsis.

**Figure 2.**
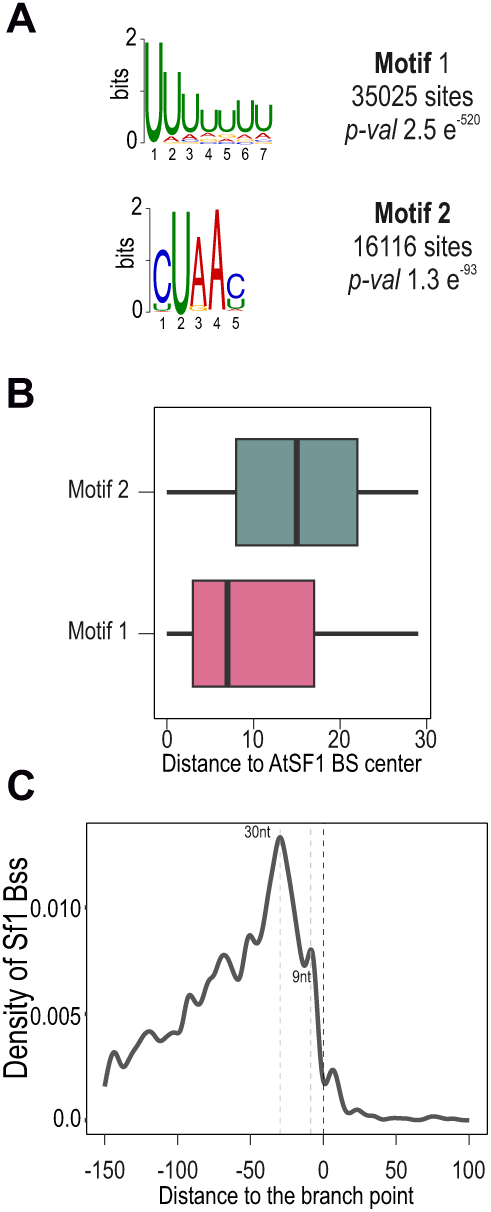
Sequence and positional features of AtSF1 binding motifs in introns. (A) Sequence logos of the most significantly enriched motifs in an area 30 nucleotides around the SF1 binding sites in introns as determined by STREME (94) (B) Positional map of the density of the two motifs shown in (A) from the center of each identified binding site to focus the comparison on the relative position of the identified motifs. (C) Density of SF1 BS centered at the branch points determined by lariat sequencing (11)

### AtSF1 predominantly affects alternative splicing by direct binding

To determine how AtSF1 influences splicing, we integrated RNA-seq from wild type and *sf1-4* plants with iCLIP binding data. Used ASpli (55), we detected 4,196 differential alternative splicing (DAS) events across 3,101 genes—hereafter referred to as differentially spliced genes (DSGs) in the *sf1-4* mutant (Table S4). Notably, ∼50% of these DSGs are direct AtSF1 targets identified by iCLIP, underlining AtSF1’s substantial, direct role in splicing.

Among splicing types, intron retention (IR) was the most frequent (∼55%), followed by alternative 3’splice site (Alt3ss), exon skipping (ES), and alternative 5’splice site (Alt5ss) events (Figure 3A). In *sf1-4* mutant, nearly 90% of the retained introns showed increased IR, reflecting widespread loss of splicing efficiency (Figure 3B). Conversely, in 75% of the altered ES events, mutations led to enhanced exon skipping, resulting in shorter mRNAs isoforms (Figure 3B), suggesting that AtSF1 normally acts as a positive regulator of exon inclusion, potentially by facilitating the recognition of splice sites. For Alt3ss events, ∼85% showed higher percent-spliced-in (PSI) values (see material and methods) in *sf1-4* compared to wild type plants, indicating selection of longer exons (Figure 3B).

**Figure 3.**
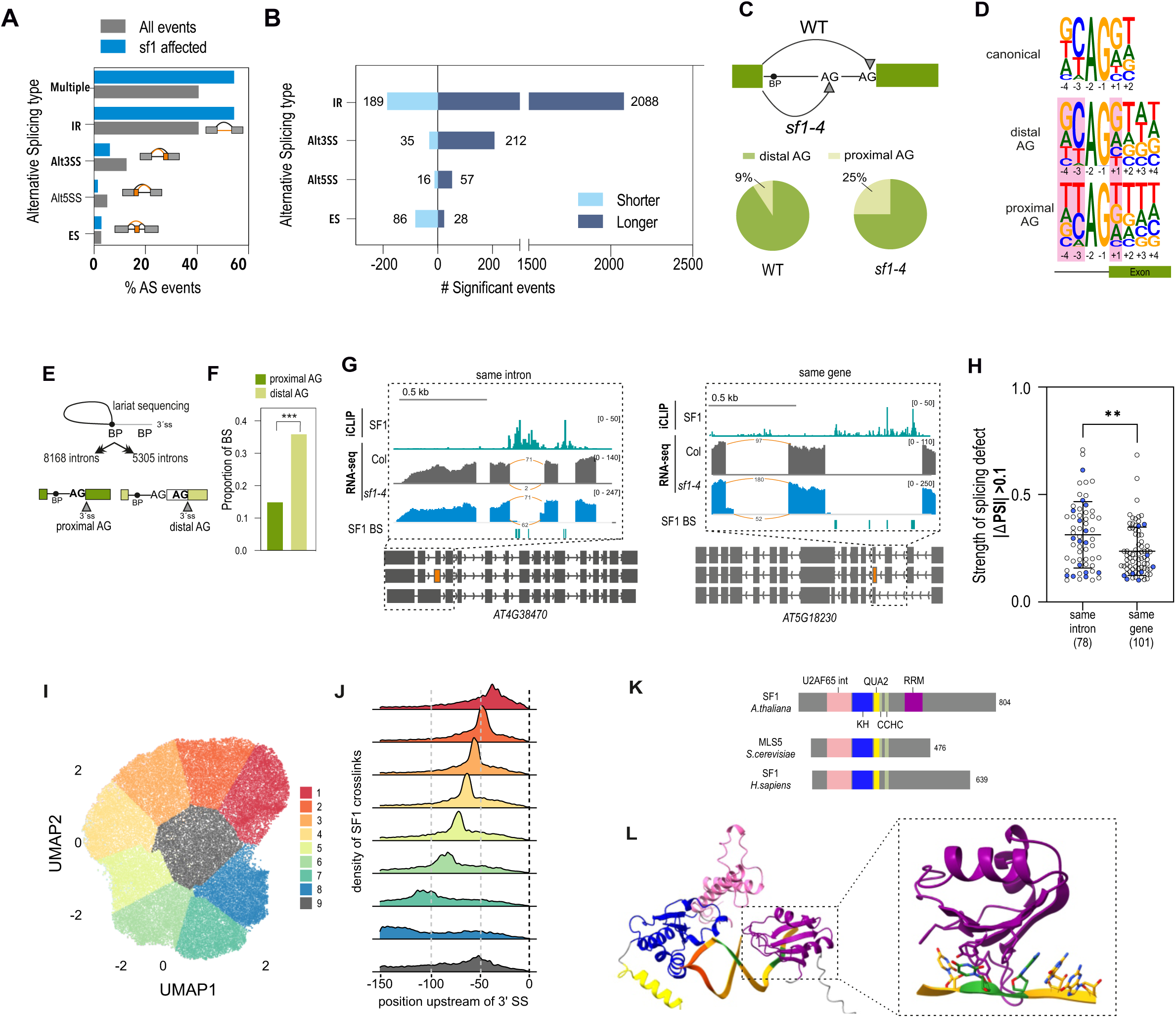
Impact of AtSF1 binding on alternative splicing regulation in Arabidopsis. (A) Relative frequencies of different types of splicing events. Grey bars refer to all expressed AS events annotated in RTD3 (46), and cyan bars show differentially spliced events between wild type and *sf1-4* mutant determined by ASpli (FDR <0.15; |ΔPSI| > 0.05). Alt3SS: alternative 3’splice site, Alt5SS: alternative 5’splice site; ES: exon skipping; IR: intron retention; Multiple: more than one type of event observed. (B) Distribution of AS events separated by type and the effect of the *sf1-4* mutation on the transcript. |ΔPSI| > 0 indicates a longer transcript (dark blue) while |ΔPSI| <0 indicates a shorter transcript (light blue). (C) Top. Schematic view of the 3’SSusage. The preferred acceptor site for each genotype is marked. Below. Pie chart showing the proportion of usage of different acceptor sites among the introns that show differential alternative 3’SSbetween wild type and *sf1-4.* (D) Pictogram of the frequency distribution of nucleotides around the proximal or distal AG. Differences in frequencies are highlighted with a red background (E) Classification of introns in two categories: introns where the first AG of the intron is used as acceptor site (proximal), or intron where the first AG is skipped and a more distal AG is used as acceptor site (distal) (F) Proportion of the SF1 intronic binding sites in the two intron categories defined in D. Fischer test was done. (G) Genome browser tracks of RNA-Seq and iCLIP for two genes to represent the two categories of events: “same intron”: where binding and splicing alteration colocalize in the same intron, or “same gene”: where binding and splicing alteration take place on a different intron. (H) The effect on splicing alteration is larger in introns that are bound by SF1 (same intron) than in introns that are not bound by SF1 (same gene). The number of events in each case is shown in parenthesis. (I) 2D uniform manifold approximation and projection (UMAP) from the top 50k crosslinked regions -150 nt to 0 nt upstream from the 3’ SS followed by k-means clustering. Clusters were renamed according to the position of the peak in the cluster. (J) Ridge-line plot displays smoothed AtSF1 crosslink densities of each cluster within the nine k-means clusters. (K) Schematic representation of domain structure of branch point proteins: SF1 from *A. thaliana*, SF1 from *H. sapiens* (Q15637) and Mls5 from *S. cerevisiae* (Q12186). Domains are highlighted in different colours. Pink: U2AF65 interaction domain; blue: KH domain; yellow: QUA2; green: CCHH knuckle domain; violet: RRM domain. Number of amino acids are shown beside each protein. (L) AlphaFold 3.0 model of the interaction between AtSF1 and a 24 nucleotide RNA: UUUAUU<u>CUAAC</u>GG**AG**UAAAUU**AG**G. The branch point sequence is underlined, splice site AG are in bold. The protein domains correspond to the colours in (K).

Sequence analysis revealed that wild type plants prefer distal AG acceptor sites (∼90% of cases), whereas *sf1-4* mutant plants shift to the proximal AG usage (Figure 3C). Splice-site strength is mostly affected by intronic rather than exonic sequences, in particular the hexamer downstream or upstream the donor or acceptor sites, respectively (56). Loss of AtSF1 drives selection of sites characterized by reduced consensus nucleotides, especially a lower occurrence of cytosine at position -3 and of guanine at positions -4 and +1 relative to the AG (9) (Figure 3D, Figure S4), indicating reduced splice site strength, reinforcing the role of AtSF1 in guiding accurate 3’ss selection through recognition of sequence features associated with strong acceptor sites.

The process of branch point selection determines 3’ss recognition, where the first AG downstream of the branch point is usually used as the 3’acceptor site (57, 58). Analysis of lariat-sequencing from 13,873 introns (11) showed that in 8168 introns the first AG after the branch point is used as 3’ss (hereafter proximal AG), whereas in 5,305 introns a more distal AG is used (distal AG) (Figure 3F). The density of BS occupancy per intron does not differ among these two groups (Figure S5). However, we found that only 15% of proximal AG introns harbor AtSF1 binding, compared to 37% of distal AG introns (p < 2.2e–16; Figure 3F), suggesting AtSF1 supports accurate 3′ site selection, particularly at alternative sites.

To test whether AtSF1 binding influences selection of the acceptor site, we analyzed the splicing effect caused by the *sf1-4* mutation across two groups of Alt3ss events: those in which AtSF1 binding occurs in the intron where splicing is affected (“same intron”), and those in which binding occurs elsewhere in the gene (“same gene”) (Figure 3G). An example of the “same intron” category is *AT4G38470*, where AtSF1 binds to intron 2, which also exhibit an Alt3ss shift (Figure 3G). In contrast, in *AT5G18230*, AtSF1 BSs are detected in introns 1 and 2, whereas the 3’ss splicing alteration is detected in intron 3 (Figure 3G). Among the 235 genes showing altered 3’ss usage, 167 harbour AtSF1 binding sites, and 78 of these fall into the “same intron” group. Collectively, comparing loci with AtSF1 binding in the affected intron versus elsewhere in the gene demonstrated significantly larger splicing effects (ΔPSI) when binding is local (“same intron”), reinforcing the mechanistic importance of direct AtSF1-RNA contact in modulating splice site choice (Figure 3G–H). We next compared the density of iCLIP crosslink sites at the altered Alt3ss events with those at non-differentially spliced annotated Alt3ss events. As expected, AtSF1 binding was depleted over exonic regions (Figure S6A). Interestingly, we observed a significant increase in AtSF1 crosslink sites upstream of the 3’ss, a region consistent with branch point proximity (Figure S6A). We extended this analysis to all the types of events and found that AtSF1 binding also correlated with Alt5ss regulation (Figure S6B), possibly by interacting with U1 snRNP components. However, ES and IR regulated by AtSF1 are not associated to AtSF1 binding (Figure S6C-D).

To address AtSF1 binding properties in relation with the acceptor site selection, we analyzed the crosslink signals up to 150 upstream of the acceptor site of the intron. Dimensionality reduction UMAP (Unsupervised uniform Manifold Approximation and Projection) (Figure S7) followed by k-means clustering of crosslink signals identified nine distinct binding profiles relative to the 3’ss (Figure 3I). Interestingly, all clusters are characterized by being mostly unimodal and having a particular peak distance to the 3’ss (Figure 3J). Alt3ss events mapped to clusters proximal to the branch point (cluster 2), while IR events localized toward intron centers (Cluster 5) (Figure S8). Whether the distance determines the fate of the splicing reaction remains to be determined.

Structurally, AtSF1 diverges from yeast and human homologs by containing a unique plant-specific RRM domain (19) in addition to conserved KH and zinc finger domains (Figure 3K, Figure S9). We performed AlphaFold3 modelling of AtSF1 and its interaction with a 24-nucleotide RNA containing the BPS defined by our iCLIP data, positioned upstream of two AG acceptor sites separated by six nucleotides. Modelling revealed high-confident interactions (SF1+RNA: iPTM = 0.86, pTM = 0.46) between AsSF1 and the RNA, with the KH domain engaging the BPS similar to what was found for yeast (12), and the RRM domain contacting RNA 3’ regions (Figure 3K). Examination of atomic distances between the RRM residues and the RNA further supports direct RRM-RNA contacts (Figure S10), even stretching the RNA molecule from its canonical conformation to maintain the interaction (Figure S11), suggesting a role in the RNA recognition mechanism of AtSF1. These data support a division of labor where AtSF1’s conserved domains mediate branch point recognition, while the plant-specific RRM facilitates downstream intron interactions, enhancing the specificity and impact of AtSF1-mediated splicing regulation in planta.

### *AtSF1* deficiency reshapes isoform architecture with functional transcript consequences

Given the widespread splicing alterations in *sf1* mutants (Figure 3A-B), we next compared gene and isoform usage to identify functionally relevant isoform switches. Differential gene expression analysis between wild type and *sf1-4* revealed 4,693 known and 259 newly annotated transcripts (46) significantly altered (FDR < 0.05; Supplemental Table S3, Figure 4A). Among genes with multiple isoforms, a substantial subset exhibited differential transcript usage: 2,234 genes switched between 4,942 isoforms (Supplemental Table S4). While there was a significant overlap between differentially expressed genes (DEGs) and those transcripts with differential isoform usage (*P* <10^-21^, Fisheŕs exact test), 1,562 genes underwent isoform switching independently of overall gene expression changes (Figure 4A). Notably, RNA-binding proteins—especially those containing RRM domains—were overrepresented among all transcripts that suffer differential isoform usage (54 genes, *P=* 2,46 e-5, Benjamini-Hochberg test). Many of these isoform switches lead to consequences affecting the transcript’s protein-coding potential (Figure 4B): the most frequent event included intron retention, disruption or loss of protein domains and changes in nonsense-mediated decay (NMD) status. For instance, 49 genes were predicted to lose a protein domain, while 72 transcripts showed altered open reading frame (ORF) length, mostly often producing shorter ORFs in the isoforms upregulated in *sf-4* (Figure 4B-C).

**Figure 4.**
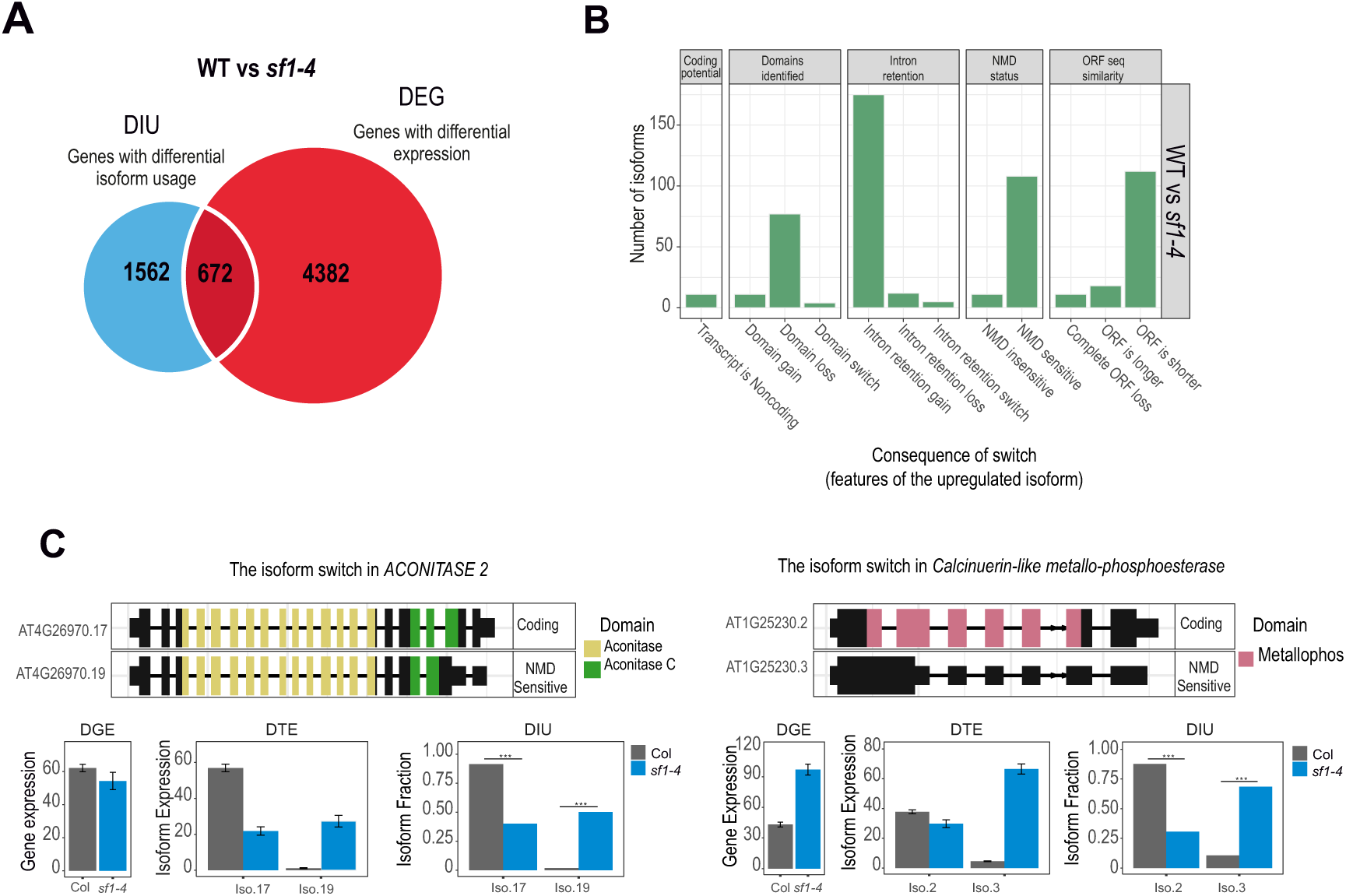
AtSF1 regulates isoform usage and transcript diversity. (A) Venn diagram depicting the extent of overlap for genes exhibiting differential isoform usage (DIU) and differential expression (DGE) identified in *sf1-4* comparison to wild-type plants (FDR < 0.05). (B) Consequences of isoform switches between wild type and *sf1-4* determined by IsoformSwitchAnalyzeR Package. Each type of switch is counted. (C) Two examples of isoform switching observed in *ACONITASE2* and *CALCINEURIN-LIKE METALLO-PHOSPHOESTERASE SUPERFAMILY PROTEIN* genes. In each case, the upper panel depicts the gene structure of the isoforms. The lower panels show differential gene expression (DGE), differential transcript expression (DTE) and differential isoform usage (DIU). Only the relevant isoforms are shown for clarity.

Specific cases highlight these functional consequences. In *ACONITASE2,* the protein-coding isoform *AT4G26970.17* is downregulated in *sf1-4,* whereas the truncated, likely NMD-targeted isoform *AT4G26970.18* is upregulated, shifting the balance toward non-functional transcripts (Figure 4C). Similarly, for the Calcineurin-like metallo-phosphoesterase gene, the non-functional isoform *AT1G25230.3* which lacks the metallosphophoesterase domain, is preferentially expressed in the mutant, reducing the proportion of transcripts encoding a functional domain (Figure 4C). Overall, isoform changes in *sf1-4* most frequently increase transcripts predicted to undergo NMD (60 vs 6 isoforms, Supplemental Table S4). Importantly, *AtSF1* deficiency does not affect the usage of alternative transcription start or termination site (Supplemental Table S4), indicating that the observed isoform diversity in *sf1-4* arises specifically from AS regulation.

### Cross-regulation of splicing factors

Approximately half of DSGs in *sf1-4* are direct AtSF1 RNA targets, compared to only 28% of DEGs (Figure 5A), demonstrating that AtSF1 predominantly regulates splicing rather than transcript abundance. Consistent with this, a pronounced upregulation of splicing factor genes was observed among the 260 splicing related factors annotated by Wang *et al.* (59), despite an overall balance of up- and downregulated transcripts (Figure 5B, Supplementary Table S3). Of the 38 splicing factor genes that were differentially expressed in *sf1-4*, 32 were upregulated (Figure 5B), suggesting a compensatory response aimed at buffering *AtSF1* deficiency and maintaining splicing homeostasis. AtSF1 binds directly 115 of these genes, and 42 also exhibited altered splicing in *sf1-4* (Supplemental Table S3, Figure 5C). Notably, this regulatory breadth surpasses human systems, where perturbation of a single splicing regulator typically affects ∼24 spliceosomal factors (60).

**Figure 5.**
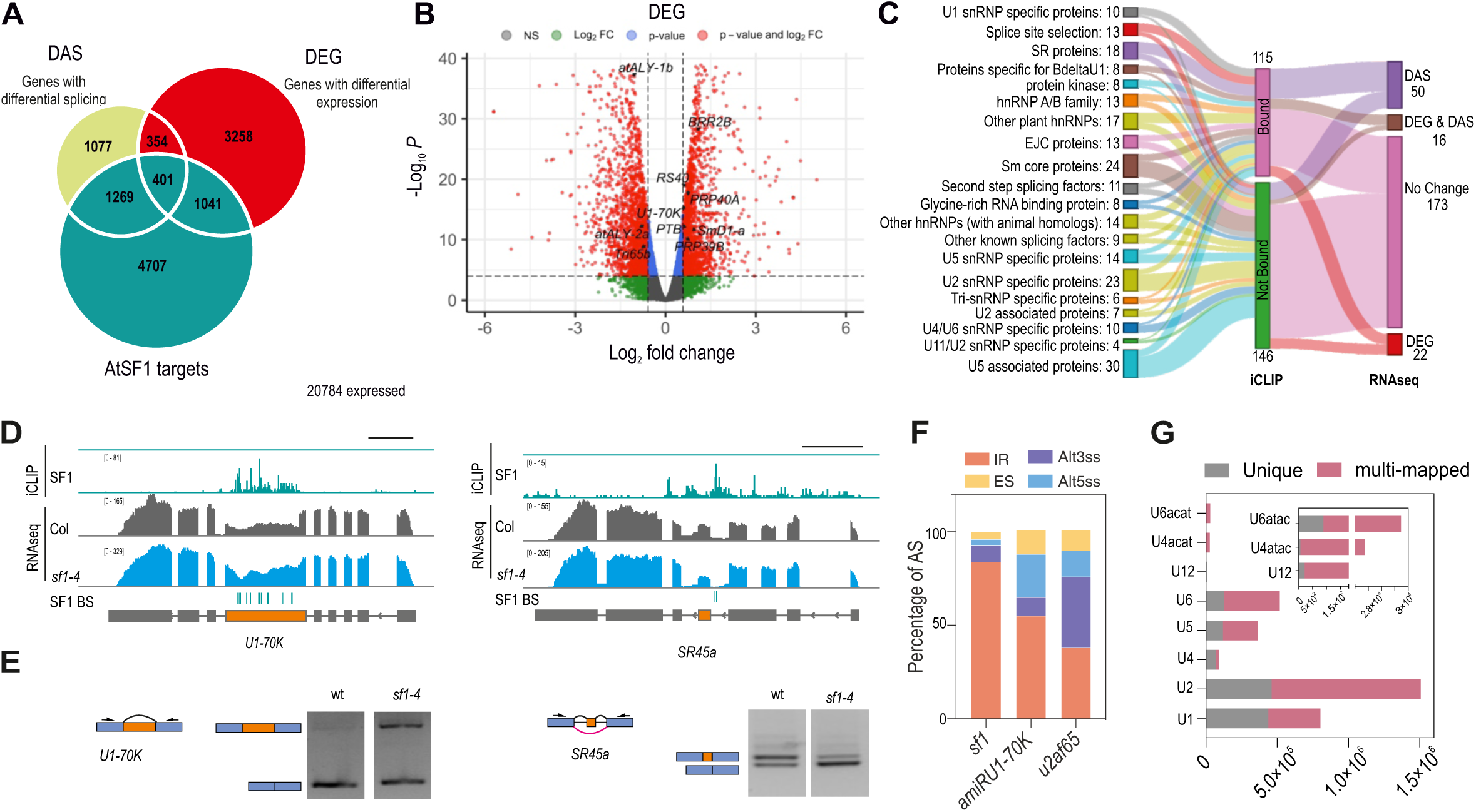
Integration of AtSF1 binding, splicing regulation, and gene expression changes. (A) Venn diagram of overlap of genes for genes differentially expressed in *sf1-4* relative to wild type (DEGs), differentially spliced genes (DSGs) and genes directly bound by AtSF1 (AtSF1 targets). (B) Volcano plot illustrating the log_2_ fold-change (x-axis) and the statistical significance as -log_10_ *P*-value (y-axis) of the RNAseq experiment for differential gene expression. The dashed line indicates the threshold above which transcripts are considered significantly differentially expressed (*P-*value <0.05). Highlighted are genes involved in splicing. (C) Sankey diagram illustrating the modes of regulation of splicing factors described in the dataset of SFs (59), indicating the number of splicing factors that are direct AtSF1 targets and the effect on the transcript when losing functional *AtSF1* (alternative splicing (DAS) or differentially expressed (DEG)). The diagram provides a visual representation that traces the flow of binding to their consequential effect on transcripts. (D) Genome browser tracks of RNASeq and iCLIP for two splicing factors, *U1-70K* and *SR45a*. Top (green). Read density of the iCLIP experiments. Middle and bottom (grey and blue) Read densities of wild type and *sf1-4* respectively. The green lines below the read densities correspond to the AtSF1 binding sites determined by iCLIP. Gene models according to RTD3 are depicted below. The alternative event is highlighted in orange. (E) Validation by RT-PCR of splicing events shown in D. Alternative regions are highlighted in orange next to the gels and position of the primers are depicted. (F) Proportion of differentially alternative spliced events in *sf1-4* (this study), *amiR-U1-70K* lines (61) and *u2af65b* (62) mutants. Available raw data was downloaded and analysed with ASpli (see methods for parameters). IR: Intron retention, Alt3ss: alternative 3’splice site, Alt5ss: alternative 5’splice site and ES: exon skipping. (G) AtSF1 binding to snRNAs. For the analysis reads were remapped to the snRNAs sequences (see methods). Unique mapping (grey) or multi-mapping (pink) is discriminated for each snRNA type. Components of the minor spliceosome are zoomed for a better evaluation.

Among the spliceosomal gene targets with altered splicing, we identified *SERINE-ARGININE RICH PROTEIN 30* (*SR30*), *ARGININE-SERINERICH SPLICING FACTOR 35* (*RS40*), *ARGININE-SERINE-RICH ZINC KNUCKLE-CONTAINING PROTEIN 32* and *33* (*RS2Z32, RS2Z33*), *SERRATE* (*SE*), *SKIP*, as well as core spliceosomal components such as *PRE-mRNA PROCESSING PROTEINS A* and *C* (*PRP40A* and *PRP40C*), *U1-70K* and *U2AF65B*. Experimental confirmation included enhanced IR in *U1-70K* transcript in *sf1-4* and the exon skipping in *SR45a* (Figure 5D-E).

To understand interactions with other spliceosome components, we compared DAS profiles in *sf1-4* with those from artificial miRNAs knock-down of *U1-70K* (61), and mutants of U2 snRNP component *U2AF65b* (62). Interestingly, significant overlap of DSGs was observed among *sf1-4*, *u2af65b*, and the *u1-70k* amiR lines, indicating regulation of common loci (Figure S12). However, overlap in specific DAS events was minimal (Figure S12), suggesting that these factors act at distinct splice sites (Figure S12). Splicing changes were consistent with the core factor roles: *u2af65b* mutants exhibited increased IR and Alt3ss usage (consistent with the role of U2AF65 in 3′ss recognition), while *amiR-U1-70K* plants exhibited IR and Alt5ss defects (Figure 5F). Limited overlap in DAS may reflect the hypomorphic nature of the alleles, as loss-of-function mutations of core spliceosome components are typically lethal.

Intriguingly, AtSF1 also bindis to major and minor spliceosomal snRNAs, including to U1 and U2, and to a lesser extent U12 and U6acat (Figure 5G). Binding to U12 snRNA occurred within the loop interacting with U11/U12-65K protein, potentially interfering with snRNP assembly. Even though U12-type introns are rare, we also observed AtSF1 binding to a subset of these introns. Of the 2,308 reported U12 introns (46), 388 were bound by AtSF1, with ∼10% of the direct targets showing mis-splicing in *sf1-4*. These findings suggest that AtSF1 also contributes—albeit modestly—to catalysing removal of U12-type introns in a subset of plant genes.

### AtSF1 contributes to circadian timing by directly binding and modulating the splicing of core clock transcripts

To identify physiological processes regulated by AtSF1, we performed a GO term analysis of DEGs (Figure 6A), DSGs (Figure 6B) in the *sf1-4* mutant, as well as on direct AtSF1 targets (Figure 6C). RNA processing and splicing emerged as dominant terms, consistent with previous findings showing broad effects on splicing factor expression in *sf1-4* (Figure 6A-B), consistent with our previous finding that splicing factors are affected in the *sf1-4* mutant (Figure 5). Notably, “Circadian regulation of gene expression” and “rhythmic process” were highly enriched amog all subsets, with the strongest enrichment found in AtSF1 direct targets of (11-fold enriched, Figure 6C).

**Figure 6.**
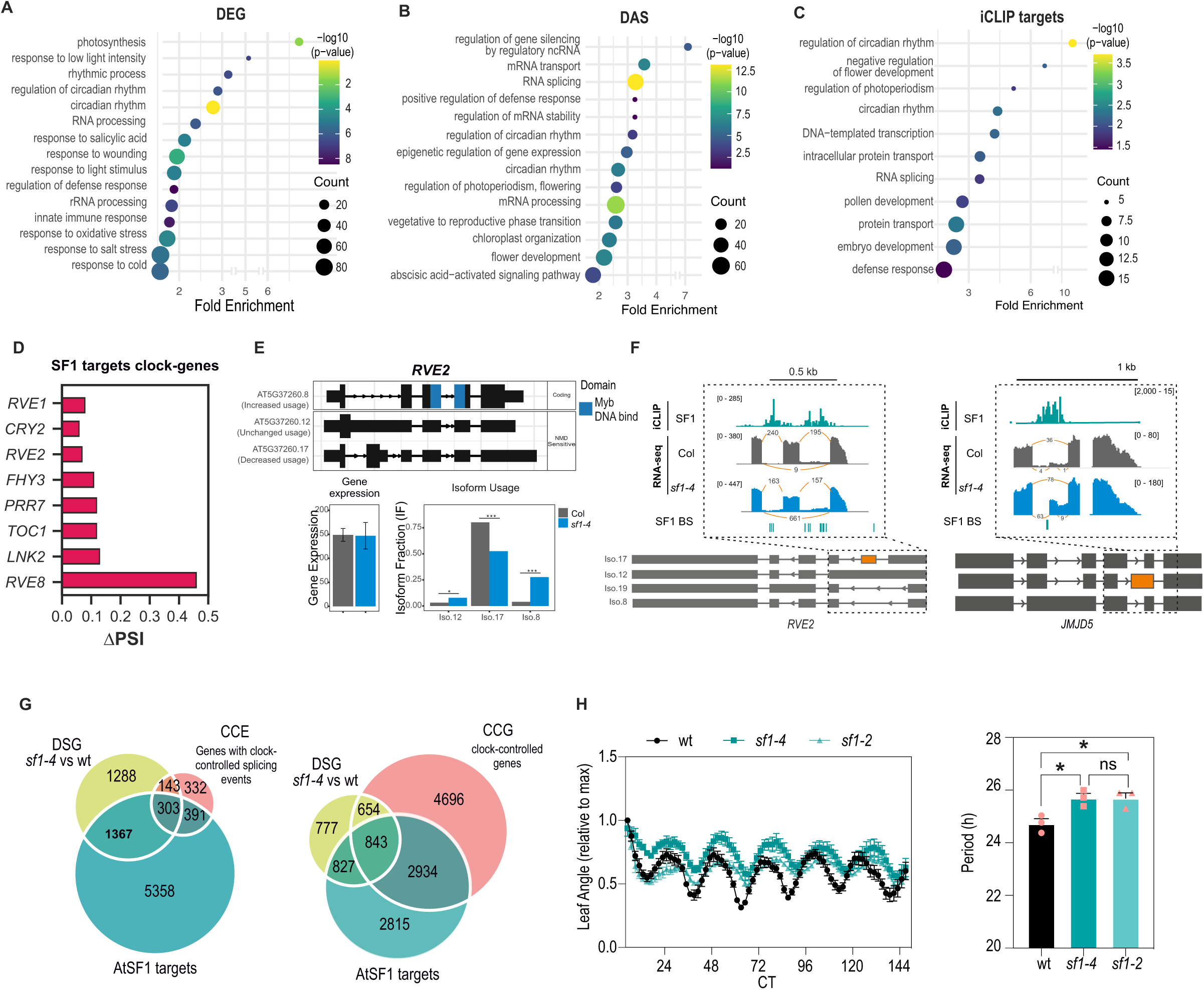
Functional implications of AtSF1-dependent splicing in circadian regulation. (A-C) GO term enrichment analysis of DEGs (A), DSGs (B) or transcripts bound by AtSF1 (C) represented as bubble plots. Gene count in each category are represented by the size of the circles. (D) ΔPSI profile of significantly differentially spliced core clock genes between wild type and *sf1-4*. (E) Isoform switching observed in *RVE2*. Total gene expression was not significantly different between wild type and *sf1-4* but significant switching was observed among the isoforms, with three isoforms exhibiting differential usage (* *P* <0.05, *** *P* <0.001). Only relevant isoforms are shown for clarity. The Myb domain is coloured in blue. (F) Genome browser tracks of RNA-Seq and iCLIP reads for *RVE2* and *JMJD5.* SF1 BSs are represented in green below the read densities. Gene models with the most important isoforms are depicted below the tracks. Alternative events are highlighted in orange. (G) Overlap between DSG and iCLIP targets determined in this study and Clock-Controlled Events (CCEs) (left) or Clock-Controlled Genes (CCGs) (right) from Romanovski et al, 2020 (63). (H) Circadian rhythm of leaf movement. (Top) Plants’ vertical leaf motion (RLM) was obtained for the first pair of leaves of seedlings entrained under long-day conditions (16 h light/8 h dark) and then transferred to continuous light (LL). WT, *sf1-4* and *sf1-2* alleles (Bottom). The period length of leaf movement rhythms is estimated by Fast Fourier transform–nonlinear least-square test (FFT–NLLS) (Right). Error bars represent SEM.

Consistent with these findings, we detected splicing alterations in multiple core clock genes, including *TOC1, REVEILLE* family members (*RVE1* and *RVE2*), *PRR5* and *PRR7, ELF3* and *NIGHT LIGHT-INDUCIBLE AND CLOCK-REGULATED GENE 2* (*LNK2*) (Figure 6D, Supplemental Table S2, S3). Several of these genes showed isoform changes with functional implications. For example, exon 2 skipping in *RVE2* is increased in *sf1-4,* elevating the level of a functional isoform (*AT5G37260.8*), that retains the MYB-DNA-binding domain (Figure 6E). This splicing event is temperature-responsive (32), and AtSF1 binding in the flanking introns suggests direct regulation (Figure 6F). AtSF1 binding was also found in other genes clock-related genes that are not part of the core oscillator, such as the histone demethylase *JUMONJI DOMAIN CONTAINING 5* (*JMJD5,* AT3G20810), where binding in intron 4 correlates with the presence of an Alt3SS event (Figure 6F). Because *jmjd5* mutants displayed changes in histone methylation, we measured global histone methylation in *sf1-4* mutants and found a modest reduction in H3K27 methylation (Figure S13). Additional clock genes such as *LNK1*, *LNK4* and *PRR3* show splicing changes independent of AtSF1 direct binding; for instance, *LNK1* displays increased exon 2 skipping in *sf1-4,* generating a truncated protein lacking the first 51 amino acids (Figure S14).

Beyond the core clock, AtSF1 regulates a broad subset of clock-controlled genes. Of 1,169 genes with clock-controlled splicing events (63), 446 are also regulated by *AtSF1,* with ∼60% of these being AtSF1 direct targets (Figure 6G). While one third of the Arabidopsis transcriptome displayed rhythmic expression (63), over 50% of the DSGs and ∼50% of the AtSF1 direct targets are classified as clock-controlled genes (63) (Figure 6G). These findings suggest a central role for AtSF1 regulating circadian rhythms in Arabidopsis. To test this hypothesis, we measured leaf movement rhythms and found that both *sf1-2* and *sf1-4* mutants has lengthened circadian period (∼1 hour) compared to wild type plants (Figure 6H).

Together, these results demonstrate that AtSF1 binds and modulates the splicing of both core clock genes and clock-controlled transcripts, thereby playing a central role in Arabidopsis circadian regulation.

### AtSF1 loss leads to widespread transcriptomic changes impacting pathogen responses

Defense-related biological processes are significantly enriched among direct AtSF1 targets as well DSGs and DEGs (Figure 6A-C). Notably, key players in salicylic acid (SA) biosynthesis and signaling during immune activation, such as *ARABIDOPSIS ISOCHORISMATE SYNTHASE* enzyme (*EDS16*), *NPR1-like protein4* (*NPR4*), *TGACG sequence-specific binding protein* 2 (*TGA2*) and the *NADPH* oxidase *RBOHD,* are downregulated in *sf1-4* (Figure 7A, Table S3), indicating broad disruption of the SA-mediated immune pathway.

**Figure 7.**
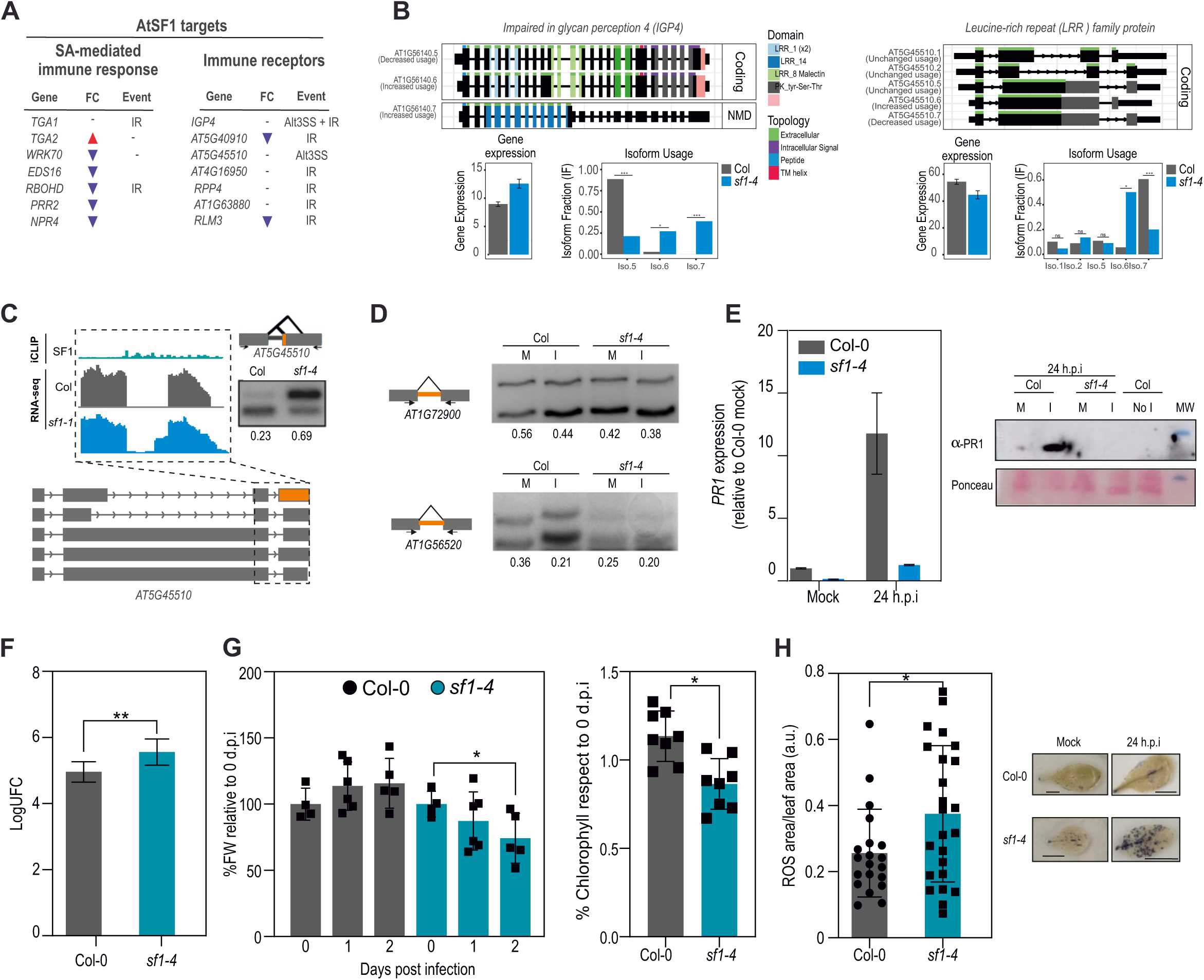
AtSF1-mediated splicing control of pathogen-related genes and immune response. (A) Table showing differential expression and differential splicing in *sf1-4* of pathogen related genes that are bound by AtSF1. Genes are dividing according to their function in plant defence. (B) Isoform switching observed for two pathogen related genes. (Top) Predicted topology and domain structure from each isoform. (Bottom) Quantitation of total gene expression and isoform usage (* *P* <0.05, *** *P* <0.001). Only relevant isoforms are shown for clarity. (C) Genome browser tracks of RNA-Seq and iCLIP reads for *AT5G45510.* RT-PCR to detect splicing defects for *AT5G45510* in wild type and *sf1-4* is shown at the top right. Scheme with alternative regions are highlighted in orange in the diagram next to the gels and the positions of the primers used are shown. The ratio of the splicing (unspliced/total) is shown below each lane. (D) Two-weeks-old plants grown in LD were infected by flood with *P. syringae* DC3000. Levels of *PR1* transcript (left) were determined by qRT-PCR and PR1 protein was assessed by western blot using anti-PR1 antibody (right) 24 h post infection in infected (I) or mock treated (M) plants. For western analysis, ponceau red staining shows equal loading across samples. (E) RT-PCR to detect splicing of *AT1G72900* and *AT1G56520* upon infection in wild type and *sf1-4* mutants. Scheme with alternative regions are highlighted in orange in the diagram next to the gels and the positions of the primers used are shown. The ratio of the splicing (unspliced/total) is shown below each lane (F) Two-weeks-old plants grown in LD conditions were infected by flood with *P. syringae* DC3000. Bacterial growth was assessed 2 d.p.i. in wild type and *sf1-4* mutant plants. Data represent the average of log-transformed bacterial growth (n = four independent biological replicates). (CFU: colony-forming units). This experiment was repeated twice with similar results. Error bars indicate SEM. P < 0.05 (one-way ANOVA followed by Tukey’s multicomparison test) (G) (Right) Fresh weight of Col-0 and *sf1-4* mutants after 0, 1 or 2 days post-inoculation with mock or *P. syringae*. (left) Chlorophyll quantification in seedlings two days post-inoculation (2 dpi), expressed relative to 0 dpi. (H) ROS quantification in leaves infiltrated at OD 0.002, measured 24 hours post-inoculation. Representative leaves are shown above each genotype.

Beyond transcriptional changes, numerous immune receptor genes- both from the pathogen-associated molecular pattern-triggered immunity (PTI) and effector-triggered immunity mediated by intracellular immune receptors of the nucleotide-binding leucine-rich repeat NLR-type, exhibit splicing defects in the mutant, including IR and Alt3ss (Figure 7A). A clear example is *AT1G56140 (IGP4),* where the functional isoform *AT1G56140.4* is markedly reduced while non-functional isoforms with retained intron or NMD-predicted premature stop codons (*AT1G56140.6, AT1G56140.7*) are increased (Figure 7B-C, Figure S15), validated by RT-PCR. Among the NLR genes, *AT5G45510* undergoes Alt3ss leading to transcripts with divergent 3’UTRs (Figure 7B-C), which may affect stability of the transcript.

We performed functional analysis of defense genes splicing from TIR-NBS-LRR family genes which are AtSF1 target genes and found that in wild type plants, infection with *Pseudomonas syringae* pv. *tomato* DC3000 using a flood inoculation assay (64) triggers IR reduction and increased splicing of genes, resulting in more functional mRNA isoforms. In non-infected wild-type plants, the ratio of unspliced to total isoforms for *AT1G72900* and *AT1G56510* were ∼0.56 and ∼0.36, respectively. Following *Pst* infection, the proportion of the spliced isoform increased relative to the unspliced transcript, suggesting elevated production of a potentially functional mRNA (Figure 7D). In *sf1-4*, these genes displayed a constitutive “infection-like” splicing profile regardless of actual infection (Figure 7D), accompanied by impaired induction of *PATHOGENESIS RELATED GENE 1* (*PR1*) expression and PR1 and PR2 protein accumulation, canonical SA-responsive markers (Figure 7E, Figure S16), consistent with a compromised SA pathway.

Phenotypically, *sf1-4* plants challenged with *P. syringae* exhibit higher bacterial titers, significant biomass loss, reduced chlorophyll, and elevated reactive oxygen species (ROS) accumulation (Figures 7F–H), indicating increased susceptibility and accelerated disease progression. These results establish that AtSF1 is critical for regulating both the transcription and splicing of immune genes; its loss renders the transcriptome constitutively mimic pathogen-challenged states, ultimately compromising effective defence signalling, although locus-specific causality remains to be established.

## Discussion

Phenotypic plasticity in plants is underpinned by transcriptome reprogramming, with AS serving as a key layer that expands proteomic diversity from single genes (65). While many plant splicing factor mutants exhibit widespread AS defects, how splicing factors engage RNA targets *in vivo* to direct biological outcomes is only beginning to be resolved. Here, by generating the first *in vivo* single-nucleotide resolution binding map of AtSF1, we unveil how branch point and 3’ss recognition by this factor mechanistically shapes AS and its downstream regulatory functions.

Our iCLIP analysis identified ∼77,000 AtSF1 BSs in 7,418 Arabidopsis genes, establishing AtSF1 as one of the most widely bound plant splicing factors to date (Figure 1). Despite the low abundance of intron-containing transcripts in total lysates, AtSF1-bound intronic sequences were successfully captured. Crosslink signals were detected throughout all introns, spanning their entire length, but significant BSs were particularly concentrated near the 3’ss (Figure 1F). Unlike other plant splicing factors studied so far (47–50), more than 60% of AtSF1 BSs mapped within introns (Figure 1E). This mirrors the pattern observed for HsSF1 (8) (Figure 1E), implying a conserved mechanism for splicing site recognition across kingdoms. Interestingly, AtSF1 exhibits a unimodal intronic binding pattern that can be grouped into different clusters that differ in their distance to the 3’ss (Figure 3I-J), contrasting with the more complex, multi-peaked binding profile reported for human SF3B1, a factor involved in later stages of splicing (66). Although crosslink peaks were largely depleted from exons and 5’UTRs, substantial AtSF1 occupancy was also observed in 3’UTRs and in intergenic regions (Figure 1). RNA polymerase II (RNAPII) intergenic transcription has been recently profiled in plants through plant Native elongating Transcript sequencing (plaNET-seq) (67, 68). We observed that most of intergenic BSs fall within these transcribed intergenic regions, which correspond either to novel transcripts or extended 3’UTRs (Figure S17). This raises the possibility that AtSF1 may participate not only in intron definition but also in 3’end mRNA processing, similar to the U1 snRNP’s proposed role in alternative polyadenylation site selection (69). Whether AtSF1 directly contributes to polyadenylation site choice remains to be determined.

Motif analysis revealed a strong enrichment for branch point-like sequences (9–11, 43) at AtSF1 BSs, yielding the first experimental evidence of AtSF1’s *in vivo* BPS specificity in plants (Figure S18). Strikingly, AtSF1 binding peaks are positioned adjacent to, rather than directly at, the branch point (Figure 2), in contrast to yeast SF1/Mls5, whose binding is sharply centered at the yeast branch point motif (UACUAAC) (70). This observation may have several explanations. First, UV crosslinking has a moderate bias towards capturing contacts at uridines (71), which could enrich signals over U-tracs and under-recover adenosine-centred branch point contacts, shifting binding peaks (Figure 2A-B). We consider this less likely, as iCLIP for HsSF1 revealed binding peaks at the BPS, with a reduced signal at the adenine from the branch point itself (8). Alternatively, the adjacent positioning could imply that AtSF1 is facilitating accessibility rather than executing branch point recognition. By binding next to the BPS, AtSF1 could stabilize the local RNA conformation, pre-bulging the branch point adenosine so that the U2 snRNA can base pair with it efficiently. In this way, AtSF1 could be considered as a positional guide rather than a definite selector of the branch point as in other organisms. Another hypothesis is that this divergence reflects the distinct domain architecture of SF1 across eukaryotes. Whereas yeast and human SF1 contain only one RNA binding domain (the KH domain), AtSF1 possessed two, both a KH and an RRM domain (Figure 3K, Figure S9). These RNA-binding modules differ markedly in both recognition mode and affinity: RRMs typically contact 2-8 nt RNAs via sequential stacking and hydrogen bonds interaction, often achieving nanomolar affinities, whereas KH domains engage shorter (∼4-5 nt) sequences primarily only through hydrogen bonding and not stacking interactions yielding micromolar affinities (reviewed in (72)). Thus, iCLIP signals in plants may reflect the RRM-mediated contacts, rather than those of lower affinity KH-mediated engagement of the branch point itself. In this model, the KH domain may still position AtSF1 at or near the BPS, while the RRM stabilizes binding to adjacent intronic regions, effectively shifting the apparent peak of occupancy away from the canonical BPS. These findings suggest an evolved, domain-specific mechanism by which plant SF1 stabilizes the local RNA environment to promote accurate 3′ss choice and robust branch point recognition.

Although splicing defects in individual genes have been previously reported in Arabidopsis *sf1* mutants (20, 40, 43, 73), comprehensive transcriptome-wide analysis of AS remained limited, and the associated datasets are not publicly available (43). Transcriptome-wide, we show that half of DSGs in *sf1-4* are direct AtSF1 targets, underlining its active and predominant role in splicing regulation. In agreement with prior studies (43), the main effect of AtSF1 loss is increased IR (Figure 3A-B), but we now demonstrate that AtSF1 binding near branch points is associated with distal 3’ss usage, securing accurate splice site selection (Figure 3, Figure 4, Figure S6A). In *sf1-4* mutant we detected a strong bias towards the use of upstream, weaker 3’ss, leading to an accumulation of the longer transcript isoforms (Figure 3, Figure 4). In some cases, these isoforms represented up to 90% of the total transcript population (Table S4), underscoring the central role of AtSF1 in safeguarding splice site fidelity. The reason why Alt3ss become more frequent in *sf1* might be explained by the fact that functional AtSF1 is not present to stabilize branch point recognition. Without anchoring close to the branch point, fidelity and 3’ss selection accuracy may drop and usage of suboptimal sites cannot be prevented. A similar splicing outcome has been recently reported for mutants in *XAP5 CIRCADIAN TIMEKEEPER* (*XTC*), that lead to aberrant 3’splice selection (74, 75). Mls5 from *Saccharomyces cerevisiae* serves to effectively splice introns with weak 5’SSor branch point mutations (76). In contrast, although human SF1 is essential for cell viability, its depletion does not alter general splicing, suggesting that it might be only required for splicing of specific pre-mRNA *in vivo* (77). In addition to its impacts on 3’ss, AtSF1 influences to a minor extent alternative 5’ss selection, possibly through regulatory effects on the U1 snRNP composition or abundance, as it binds the U1-70K transcript and alters its processing (Figure S6B). However, IR or ES events do not appear to be strongly driven by AtSF1 binding per se (Figure S6C-D). This is consistent with introns retained in *sf1-4* showing broad AtSF1 crosslinks signals across their entire length (Figure S6C), reinforcing the idea that positional specificity, as observed near 3’ss, rather than absence of binding, underlies splicing effects. Interestingly, Hs SF1 has been recently linked to the activation of exon inclusion as we observed for AtSF1 (Figure 3B). Notably, competition for branch point sequences within the intron between SF1 and Quaking protein determines the fate of the alternative exon: while SF1 promotes exon inclusion, the presence of Quaking represses inclusion by inhibiting SF1’s access to the branch point (78).

The *sf1-4* allele carries a T-DNA insertion upstream of the RRM domain. Inspection of the RNA-seq reads confirms that a truncated *AtSF1* transcript accumulates in *sf1-4* to levels similar to those in wild type plants (Figure S19), suggesting that in the mutant, a truncated protein could be produced. The predicted truncated AtSF1 protein retains the KH, QUA2 and Zinc-Finger domains but lacks the RRM domain, therefore potentially retaining sufficient activity to support viability (Figure 1A) while compromising its splicing functions (Figure 3, Figure 4). AlphaFold3 modelling indicates the truncation disrupts RNA contacts at the 3′ end without globally altering the remaining domain configuration (Figure S11), providing a structural rationale for the global shift toward proximal 3’ss usage and impaired fidelity observed in *sf1-4* (Figure 3, Figure 4). Previous observations demonstrated that the RRM domain of AtSF1 contributes to splicing regulation of a specific substrate, namely *HSFA2* (20). Taken together, these observations support the hypothesis that the mutant protein did not lead to significant differences in the branch point recognition, but the loss of the RRM domain in the *sf1-4* mutant may reduce intron binding, which could shift splicing towards the proximal AG acceptor sites and altering 3’ss selection (Figure 8). In sum, we provide mechanistic implications of AtSF1 domain architecture.

**Figure 8.**
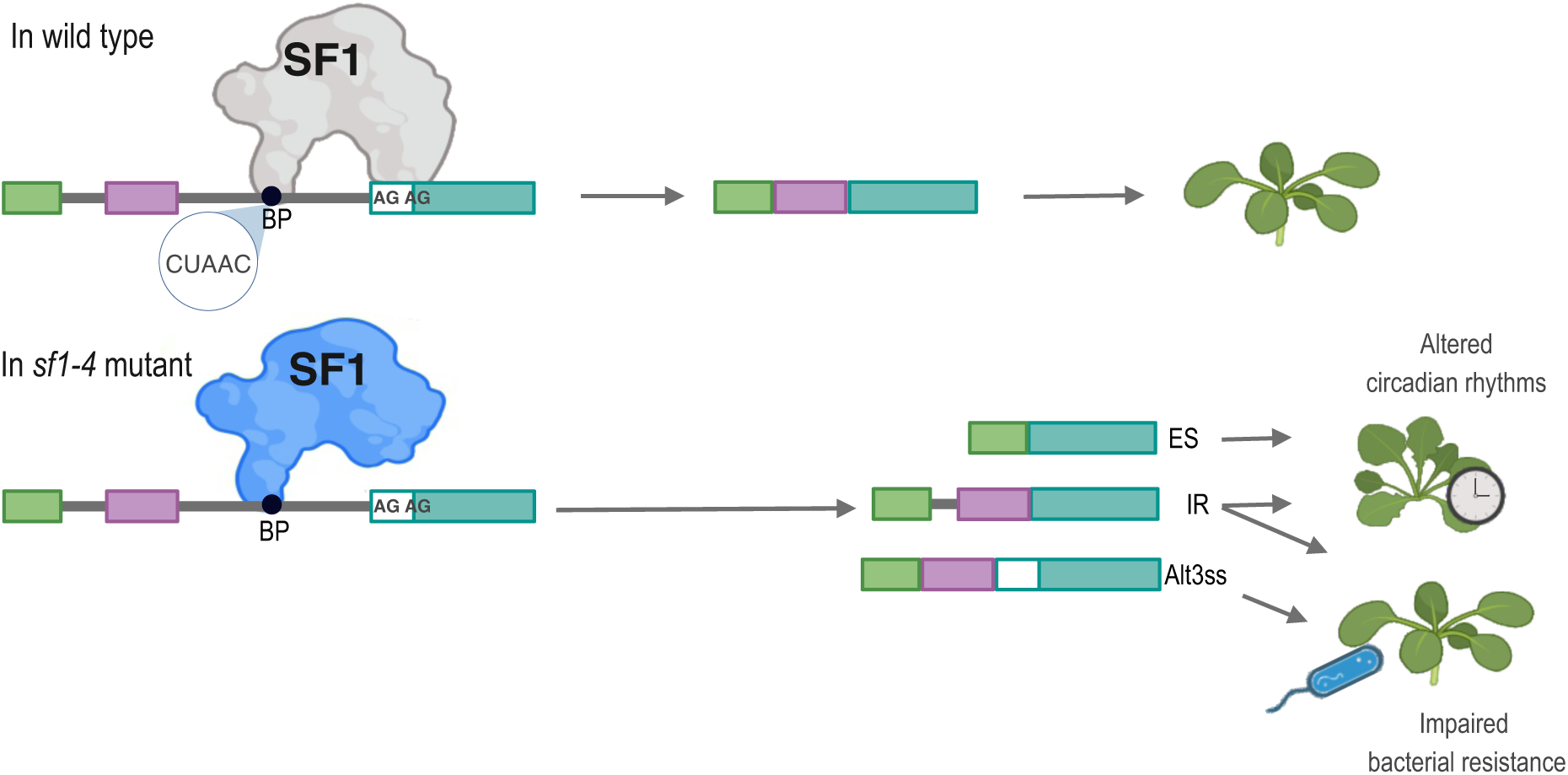
Proposed model summarizing AtSF1 functions in RNA processing and stress responses. In wild type plants, splicing occurs normally and AtSF1 uses preferentially the distal AG towards the proximal AG, leading to the shorter option of the final transcript. In *sf1-4* mutants, splicing is affected. In particular, a strong bias towards the proximal AG is observed, and the mRNA possess a shorter version of the intron rendering a longer transcript. This switch in AG usage brings along widespread changes in the transcriptome. Impact of the mutation in the phenotype is exemplified here. IR and ES events on clock genes are associated with the long period phenotype observed in the mutant, while IR and Alt3ss events on defense genes alter pathogen responses.

Functionally, AtSF1-mediated splicing is crucial for two major physiological pathways in Arabidopsis: circadian clock regulation and pathogen defense. We demonstrate that AtSF1 binds numerous core clock and clock-regulated genes (Figures 6C–G), and disruption in *sf1-4* leads to mis-splicing, altered isoform ratios, and lengthening of the circadian period (Figure 6H). AtSF1 binding often co-localizes with the affected introns (Figure 6F), supporting a direct regulatory role and implicating AtSF1 as a conduit for integrating environmental signals into the clock at the level of transcript isoform regulation. On the other hand, in the case of *TOC1* for example, AtSF1 is detected in intron 1, but IR occurs downstream of binding at intron 4 (Figure S13), an IR event previously shown to be induced by low temperature (79). Likewise, the histone demethylase gene *JMJD5* exhibits an Alt3’ss event that truncates the catalytic domain important for demethylation (80). While *jmjd5* mutants show increased methylation at H3K27 (81), *sf1-4* showed decreased levels of this mark (Figure S13). Therefore, whether this phenotype can be associated to mis-splicing of *JMJD5* in the *sf1-4* mutant remains to be determined. Downregulation of this isoform has been previously described as a light-regulated splicing switch (82) and we observed that in *sf1-4,* the splicing pattern resembles dark-grown wild type plants (Figure 6F). At least three of all these events in clock genes overlap with known temperature-sensitive splicing switches (32, 79, 81) suggesting that AtSF1 integrates environmental information into the clock machinery through isoform-specific regulation of key transcripts, thus linking splicing control with circadian adaptation.

Our data reveal that defense-associated processes are strongly enriched among direct AtSF1 targets, DSGs and DEGs (Figure 6A-C, Figure 7A). For example, the *IGP4* locus undergoes a marked shift from coding isoforms toward intron-retaining and NMD-prone Alt3ss variants, reducing the pool of functional protein and linking AtSF1 activity to immune competence (Figure 7C-D, Figure S15). Howard *et al*. (34) reported numerous transcriptome changes in response to *P. syringae* infection, including *AtSF1* which is upregulated from 1 to 6 hours post-infection (hpi) but downregulated at 12 hpi, suggesting that modulation of *AtSF1* may be part of a complex transcriptional reprogramming in the plant. We further show that pathogen infection alters the splicing of a subset of defense-related genes that are direct AtSF1 targets, and that *sf1-4* mutants constitutively display “infection-like” splicing signatures (e.g., *AT1G56520*, *AT1G72900*) even in the absence of bacteria (Figure 7D). This premature switch in isoform balance likely generates a transcriptome that is primed but not functionally protective, consistent with the elevated bacterial titers, excessive ROS accumulation, and biomass loss observed following infection (Figure 7D, F-H), despite the absence of canonical PR gene activation (Figure 7E). These findings parallel regulatory paradigms such as *RESISTANCE TO PSEUDOMONAS SYRINGAE4* (RPS4), where immune output depends not simply on the presence of multiple isoforms but on their dynamic, stimulus-responsive ratios (35) Moreover, because *EDS16* and other SA-pathway factors are mis-spliced in *sf1-4* (Figure 7A), defects in SA signaling may also contribute to the susceptibility phenotype. These changes are, however, most plausibly a downstream consequence of AtSF1-dependent splicing dysregulation rather than an intrinsic lesion in SA metabolism. In line with our finding, other splicing related proteins have been previously associated to plant immunity (21, 36, 83–86) and we have shown that a tight posttranslational regulation of splicing factors can adjust their biological activity in defense and other stresses (21). Together, these data support a model in which AtSF1 safeguards temporal specificity in plant immunity by ensuring that defense-associated splicing programs are activated only upon pathogen perception rather than constitutively deployed in basal conditions.

PTI forms a core defense layer that restricts pathogen growth, and mounting evidence indicates that its timing is coordinated by the circadian clock (87). Pathogens can exploit this connection by perturbing clock function to enhance susceptibility, in part through nuclear pore disruption. Consistent with this, we found AtSF1 binding at *NUP205*, a pore complex component previously shown to be linking clock function with immune response (87), and *sf1-4* mutants have increased intron retention (Figure S20). Together, these observations suggest that AtSF1 preserves immune competence not only by regulating defence-gene splicing, but also by maintaining functional expression of clock–immune integrators such as NUP205.

Collectively, these findings position AtSF1 as a molecular gatekeeper that ensures the temporal and contextual production of functional isoforms in key signaling pathways. When AtSF1 function is compromised, non-productive transcripts accumulate, undermining immune competence and disrupting circadian timing. More broadly, the unique RNA-binding domain architecture of plant SF1—distinct from fungal and metazoan homologs— enables engagement with multiple intronic features confering additional regulatory plasticity, while Arabidopsis *sf1* lines provide a powerful system to dissect spliceosome mechanics apart from lethality.

In conclusion, AtSF1 orchestrates branch point recognition and splice-site selection in vivo, coupling alternative splicing to critical outputs in plant development and stress adaptation. Our results establish AtSF1 as an integrator of post-transcriptional control with physiological signalling, offering new insight into the evolution of splicing mechanisms and the diversification of regulatory capacity in plants.

## Materials and methods

### Plant material and growth conditions

All *Arabidopsis thaliana* mutants are in the Columbia-0 (Col-0) ecotype. The *sf1-4* (SAIL_598_A02) (43) and *sf1-2* (SALK_062177) (40) mutants were obtained from ABRC. For phenotypic characterization, plants were grown on soil at 22°C under long day cycles (16 h light/8 h dark); 80 μmol.m−2s−1 of white light. For the RNA-seq experiments, seeds were surface sterilised, sown on Murashige and Skoog (MS) medium containing 0.8% (w/v) agar and stratified for 3 d in darkness at 4°C. Seedlings were grown in a growth chamber under controlled conditions in continuous light for 12 days.

### Generation of SF1:GFP transgenic lines

The coding sequence of *SF1* was amplified by PCR using primers described in Table S7 and cloned into pDONR/Kan (Invitrogen) via the Gateway method. For overexpressor lines with GFP N-terminal tag, the pEntry vector was recombined to pB7WGF2. The binary vector was introduced into *Agrobacterium tumefaciens* GV3101 strain and transformed into *sf1-4* plants using the floral dip method (88). Transgenic lines were selected on MS medium supplemented with 15 mg/L phosphinotricin (Duchefa).

### Immunoblots and methylation status

Plants were grown according to what is described for each experiment. Total proteins were extracted in extraction buffer (50 mM Tris–HCl pH 7.2, 100 mM NaCl, 10% (v/v) glycerol, 0.1% (v/v) Tween-20, 1 mM phenylmethylsulfonyl fluoride (PMSF), and plant protease inhibitor (Sigma)) and then centrifuged at 12,000 × g and 4 °C for 15 min. Equal amounts of protein (36 µg for PR and 100 µg for histone methylation levels) were loaded onto SDS–PAGE, quantified by Bradford Protein assay, electro-transferred onto nitrocellulose membranes and probed with anti-PR1 (Dilution 1/1,000) (Agrisera, AS10 687), anti-PR2 (Dilution 1/1,000) (Agrisera, AS12 2366), anti-H3K27me2 (Diagenode, pAb-046-050), anti-H3 (Diagenode, C15200011) antibodies overnight followed by secondary antirabbit antibody coupled to peroxidase (Dilution 1/5,000) (Invitrogen, 31460). Proteins were visualized using the ECL kit (Pierce, USA) in an ImageQuant LAS 4000, GE Healthcare. Ponceau staining is shown as loading control.

### Circadian Leaf Movement Analysis

For leaf movement analysis, seeds were sown on soil and entrained in LD (16 h light/8 h dark) conditions at 22^◦^C for seven days until the first pair of leaves were fully expanded. Then, plants were transferred to continuous light (LL) and constant temperature (22^◦^C). Photos were taken every hour for six days and analysed by recording the plants’ vertical leaf motion (relative vertical motion: RLM) with a program developed in java (89) based on the Matlab code from the software TRiP: Tracking Rhythms in Plants (https://github.com/KTgreenham/TRiP) (90). The circadian period was obtained via fast Fourier transform nonlinear least-squares (FFT-NLLS) analysis using the online program BioDare2 (91). The first 24 h were excluded from the analysis to remove potential noise caused by the transfer from entrainment conditions to constant conditions.

### RNA extraction and quantitative real-time PCR

Total RNA was isolated from seedlings using TRIZOL (Ambion) and treated with RNase-free DNase I (Promega) to remove residual genomic DNA. One microgram of total RNA was used for reverse transcription using SuperScript II reverse transcriptase (Invitrogen). Transcript levels were quantified by quantitative PCR in a Stratagene MX3005P instrument (Agilent Technologies) using *PP2A* (*AT1G13320*) or *IPP2* (*AT3G02780*) as the housekeeping gene. The sequences of the primers used to quantify expression are listed in Table S7.

### Validation of splicing events

cDNAs were synthesised as above but with SuperScript III reverse transcriptase (Invitrogen). PCR amplification was performed using SuperFidelity Taq Enzyme (Invitrogen). Primers used for amplification are detailed in Table S7. RT-PCR products were incubated with SYBR Green before electrophoresis on 2.5% (w/v) agarose gels. We selected the events to be validated according to Agrofoglio et al, 2024 (21).

### RNA-seq analysis

For RNA-seq experiments, wild type and *sf1-4* seedlings were grown on MS medium containing 0.8% (w/v) agar for 12 days under continuous light at 22°C. Three biological replicates were collected, whole seedlings were harvested and total RNA was extracted with RNeasy Plant Mini Kit (QIAGEN) following the manufacturer’s protocols. To estimate the concentration and quality of samples, NanoDrop 2000c (Thermo Scientific) and the Agilent 2100 Bioanalyzer (Agilent Technologies) with the Agilent RNA 6000 Nano Kit were used, respectively. Libraries were prepared following the TruSeq RNA Sample Preparation Guide (Illumina). Briefly, 3 μg of total RNA was polyA-purified and fragmented, first-strand cDNA synthesized by reverse transcriptase (SuperScript II, Invitrogen) using random hexamers. This was followed by RNA degradation and second-strand cDNA synthesis. End repair process and addition of a single A nucleotide to the 3′ ends allowed ligation of multiple indexing adapters. Then, an enrichment step of 12 cycles of PCR was performed. Library validation included size and purity assessment with the Agilent 2100 Bioanalyzer and the Agilent DNA1000 kit (Agilent Technologies). Samples were pooled to create six multiplexed DNA libraries, which were pair-end sequenced with an Illumina HiSeq 1500 at INDEAR Argentina, providing 100-bp pair-end reads. Three replicates for each genotype were sequenced.

### Alternative splicing, differential expression and isoform switch analysis

Alternative splicing (AS) analysis was performed as previously described in Agrofoglio *et al.*, 2024 using R with the ASpli package (Mancini *et al.*, 2021). For expression analysis, genes with a false-discovery rate (FDR) <0.05 and FC >|1.5|) were considered as differentially expressed (DEGs) between genotypes. For splicing analysis, PSI (percent of inclusion) and PIR (percent of IR) were calculated and differential splicing was considered for bins with FDR <0.15 and ΔPSI/PIR >0.05. For the analyses of RNA-seq data from artificial *miR-U1-70K* lines from (61) and *u2af-65a* mutants (62) bam files were downloaded from EBI and treated as described above to compare splicing events. For isoform switch analysis IsoformSwitchAnalyzeR Package was used (92).

### Individual-nucleotide-resolution UV cross-linking and immunoprecipitation (iCLIP) and bioinformatic analysis of iCLIP

iCLIP was performed as recently described by Lewinski et al, 2024 (41) with slight changes. Plants expressing GFP:SF1 under the 35S CaMV promoter were grown in MS plates for 12 days in continuous light. 8 g of tissue were used for each replicate. 35S:GFP plants were used as negative control. Equal amount of plant material was used and all replicates were grown in parallel. We performed 4 independent biological replicates for SF1 and 3 replicates for the negative control. Plants were crosslinked with 4000 mJ/cm^2^ UV light using the UVP CL-3000 UV crosslinker. Extracts were incubated with GFP-Trap^®^ agarose beads (Proteintech) for two hours and processed as described (41). For preparative libraries for SF1 samples, 16, 17, 15 and 15 cycles in total were used for replicate 1, 2, 3 and 4 respectively. For the GFP controls, 17 cycles were used for each replicate. Sequencing was performed at the Genomics Core Facility at the Institute of Molecular Biology (Mainz, Germany), single-end 150 nt reads.

### iCLIP data analysis

iCLIP-seq data from plants expressing SF1-GFP and GFP only controls were processed using a standardized pipeline from Lewinski et al. (2024). Raw Illumina reads were first inspected with FastQC (v0.11.9) and barcode distributions checked with GNU awk. Reads were then demultiplexed and 3′ adapters removed with Flexbar (v3.5.0), reads shorter than 15 nt discarded, and a second Flexbar pass applied for quality- and length-trimming to ensure a minimum read length of 15 nt and a good read quality. Unique molecular identifiers (UMI) were transferred to the FASTQ read-ID field and PCR duplicates later removed with umi_tools (v1.1.4) by considering UMI tags and mapping coordinates. The modified reads were aligned with STAR (v2.7.3a) to the *Arabidopsis thaliana* TAIR10 genome and the AtRTD3 transcriptome with settings that report only unique mappings and soft clipping of 3′ ends to preserve the crosslink location. For the U snRNA analyses a separate STAR index was generated from U snRNA sequences (`--runMode genomeGenerate --genomeSAindexNbases 5`) and mapping allowed multimappers up to 30 locations (`--outFilterMultimapNmax 30 --outMultimapperOrder Random --runRNGseed 42 --outSAMmultNmax 1`) assigning multimapped reads randomly. Uniquely mapped, deduplicated reads from each replicate were merged and peaks called separately for SF1-GFP and GFP only samples with PureCLIP (v1.3.1). The 1-nt resolution peaks and crosslink tracks of each replicate were then used for binding site definition and refinement using the R/Bioconductor package BindingSiteFinder (DOI: 10.18129/B9.bioc.BindingSiteFinder). By applying the initial peak-score filtering we excluded called peaks which PureCLIP score were below the 10% threshold. The empirical binding site width was determined using the step wise approach comparing the signal-to-flank ratio at to the binding sites. The reproducibility cutoffs were derived from the distribution of crosslink counts across replicates (using a 20% quantile-based cutoff), and binding sites were retained when they met replicate-specific crosslink thresholds in at least two out of four (SF1-GFP), and one out of three (GFP only) biological replicates. Final BSs were annotated by overlapping coordinates with TAIR10 and AtRTD3 gene models and classified into protein-coding, non-coding and intergenic. Features such as the 5′ UTRs, CDS, introns, and 3′ UTRs were specified additionally for protein-coding genes. Binding sites located within chloroplast or mitochondrial loci were excluded from further analysis.

For motif discovery, strand-specific sequences corresponding to reproducible binding sites were extended by 30 nucleotides upstream and downstream, resulting in 65-nucleotide regions, which were extracted using Bedtools and the TAIR10 genome FASTA file. These sequences were analysed for *de novo* RNA motifs using STREME (v5.5.1), with matched background sequences generated by randomly redistributing an equivalent number of binding sites across the target transcript over 100 iterations. Crosslink positions near splice junctions and their relationship to sequence motifs were evaluated using R and Bioconductor packages (GenomicRanges, rtracklayer) and tested for positional enrichment using nonparametric comparisons (Wilcox rank test) in fixed-size bins (for example 10-nt bins across −100 to +100 nt around junctions). visualizations and summary tracks for genome browser inspection were generated from uniquely mapped read coordinates using Bedtools (2.30) and bedGraphToBigWig (https://www.encodeproject.org/software/bedgraphtobigwig/).

### UMAP and clustering of SF1 crosslinks upstream of 3’ SS

To classify distinct SF1–GFP binding patterns upstream of 3’ splice sites (3’ SS), we defined bound regions by intersecting iCLIP2 crosslink sites with intronic sequences extending up to 150 nucleotides upstream of annotated 3’ SS, based on representative gene models. This procedure identified 99,848 intronic regions containing at least one SF1–GFP crosslink, comprising a total of 9,678,795 crosslinks distributed across 2,814,703 unique crosslink sites.

For downstream analysis, we constructed a crosslink count matrix from the 50,000 most highly covered regions, considering crosslinks within the interval from –150 to 0 relative to the 3′SS. Each region was normalized individually using the `density()` function in R to account for differences in coverage.

The resulting scaled crosslinking matrix was subjected to dimensionality reduction using UMAP ((93); https://cran.r-project.org/package=umap) with the following standard-deviated parameters:

▪ `n_neighbors = 50`
▪ `min_dist = 0.001`
▪ `metric = “euclidean”`
▪ `init = “spectral”`
▪ `n_epochs = 1000`

The UMAP embedding produced a single two-dimensional data cloud. To identify subgroups of crosslinking patterns, we applied k-means clustering, partitioning the data into nine distinct clusters of approximately equal size. Cluster numbering was subsequently adjusted manually according to the relative position of the distribution peak within each region.

### Protein and RNA Structure Prediction Using AlphaFold 3 Metrics

Computational structure predictions of the wild-type SF1 protein (Uniprot ID: Q9LU44) and predicted SF1.4 variant from *sf1-4* mutant were carried out using AlphaFold 3 (DeepMind, London, UK) through the AlphaFold server (https://alphafoldserver.com). This approach was also used to predict the complex structure involving the protein-RNA interaction.

All AlphaFold predictions were performed using default software parameters and model configurations provided by the server. The highest-ranked models were selected for downstream analysis. The pTM score is a quantitative measure that estimates the accuracy of a predicted protein or RNA structure by comparing it to a reference or native structure. It ranges from 0 to 1, with values closer to 1 indicating higher confidence and better model quality. The ipTM score similarly ranges from 0 to 1 and evaluates the confidence in the predicted interfaces within multimeric complexes, reflecting the reliability of intermolecular contacts such as protein-RNA interactions. We considered that high quality predictions had a pTM>0.5 and ipTM>0.8, following the criteria given by DeepMind.

Both AtSF1 and its mutant protein contain intrinsically disordered regions that were predicted by AlphaFold3 with low confidence. These disordered segments were not implicated in the interaction with RNA based on preliminary analyses. To improve the overall quality and confidence of the models, these disordered regions were omitted. For AtSF1-RNA interaction, we generated an artificial interacting RNA of 24 nucleotides with the following sequence UUUAUUCUAACGGAGUAAAUUAGG that includes the branching point detected in the iCLIP analysis and bibliography (CUAAC), separated by 2 nucleotides of the first AG and a distance of 6 nucleotides of the second AG. This RNA provides high pTM and ipTM scores with wild-type AtSF1 and the AtSF1.4 variant.

### Protein and RNA structure visualization using ChimeraX

Molecular graphics and analyses were performed with UCSF ChimeraX v1.10, developed by the Resource for Biocomputing, Visualization, and Informatics at the University of California, San Francisco, with support from National Institutes of Health R01-GM129325 and the Office of Cyber Infrastructure and Computational Biology, National Institute of Allergy and Infectious Diseases. It was used to visualize the .cif structures provided by AlphaFold3. It was also used to measure the distances between the relevant amino acids from SF1 and the nucleotides from the interacting RNA. We measured the distances between the centroids of the aromatic chains and the angle between them, to assess possible π-π interaction. Interactions with centroid–centroid distances of 3.3–3.8 Å and angles between 0–30° were classified as parallel. Distances of 3.8–4.5 Å and angles between 60–90° were classified as T-shape/edge-to-face.

### Accession numbers

The RNA-seq raw data has been uploaded to the BioProject database and are available under BioProject ID: PRJNA1336518. Data from iCLIP experiment is deposited under the BioProject: PRJNA1356676.

## Acknowledgements

We thank Łukasz Szewc for critical discussion of the manuscript and Kristina Neudorf and Elisabeth Klemme for technical assistance. We thank Federico Arieĺs lab for an aliquot of the antibodies detecting histone marks.

## Author contriutions

J.L.M. D.S. and M.J.Y. designed the research; Y.C.A., M.J.I., M.J.de.L., E.H., and J.L.M. performed the experiments, S.T performed AlphaFold modelling, J.L.M. and M.L performed the bioinformatic analysis, J.L.M., wrote the paper with contribution of D.S, M.J.I and Y.C.A. All authors read and approved the manuscript.

## Supplementary Datasets

**Table S1. Individual-nucleotide resolution crosslinking and immunoprecipitation (iCLIP) read statistics.**

Table of read, called peak and binding site counts for each sample/replicate and processing step.

**Table S2. Coordinates of At-SF1 binding sites identified by individual-nucleotide resolution crosslinking and immunoprecipitation (iCLIP) in *Arabidopsis thaliana*** Table with genomic coordinates of GFP-AtSF1 binding sites with the corresponding PureClip scores. Identifiers of associated gene type (protein-coding, tRNA, rRNA, snoRNA, snRNA, lncRNA, ncRNA, miRNA, nontranslating_CDS, intergenic), as well as the position of the binding sites in the annotated transcript regions (5’UTR, CDS, intron, 3’ UTR) are indicated. chrom - chromosome; UTR5 and UTR3 - 5’ and 3’ untranslated regions; CDS - coding sequence.

**Table S3. List of iCLIP targets**

**Table S4. Splicing analysis. List of differential usage of events obtained from RNA-seq from *sf1-4* and wild type at 12-h light/12-h dark.** A separate list of genes affected in splicing related to splicing metabolism is included.

**Table S5. Differential expression analysis. List of differentially expressed genes obtained from RNA-seq from *sf1-4* and wild-type at 12-h light/12-h dark.**

**Table S6. Summary statistics of Differential Transcript Usage using IsoformSwitchAnalyzeR**

**Table S7. Primers used in this study**

